# Streptococcal nucleases and host-complement interactions

**DOI:** 10.64898/2026.11.26.769382

**Authors:** Isabella A. Bennig, Styliani Kostaloupi, Lotta J. Happonen

## Abstract

*Streptococcus pyogenes* and *Streptococcus pneumoniae* are major human pathogens responsible for a wide spectrum of diseases, ranging from mild infections to life-threatening conditions. Both species have evolved sophisticated strategies to evade host immunity, including the deployment of cell-wall associated or secreted nucleases that degrade neutrophil extracellular traps (NETs), hence allowing for the bacteria to evade the host innate immune defenses. While the nucleases Sda1 from *S. pyogenes* and EndA from *S. pneumoniae* have been characterized for nuclease activity and their structures have been experimentally determined, their host interactomes beyond DNA in NETs remain poorly defined. Here, we systematically mapped the human and mouse blood plasma interactomes of EndA and Sda1 using affinity-purification mass spectrometry (AP-MS). In human plasma, EndA and Sda1 bound multiple complement components including those belonging to the membrane attack complex (MAC) as well as the complement inhibitors clusterin and vitronectin. These key interactions are highly similar to those recently described for the streptococcal nuclease A, SpnA. In line with what has been described for SpnA, the mouse plasma interactomes for both nucleases studied here were sparse, with only a few specific interactors and minimal complement engagement. Furthermore, we found that EndA follows the pattern of SpnA and is a functional inhibitor of the MAC. Together, these findings reveal a previously unrecognized role for EndA and Sda1 in human immune system evasion and underscores the highly specific evolutionary adaptation of these streptococcal species to their human host.

## INTRODUCTION

The complement system provides a first line of defense and is a central part of innate immunity, functioning as an immediate and potent effector against microbial pathogens. It comprises a network of blood plasma and membrane-associated proteins that, upon activation via the classical, lectin, or alternative pathway, orchestrate a series of proteolytic events culminating in the formation of the membrane attack complex (MAC). The MAC embeds into the lipid bilayers of target cells, forming pores that disrupt membrane integrity and result in cell lysis. Although the MAC has been primarily characterized for its lytic activity against Gram-negative bacteria, accumulating evidence indicates that MAC can also assemble on the surface of Gram-positive bacteria ^1,2^. Beyond direct killing, complement activation enhances phagocytosis, recruits inflammatory cells, and bridges innate and adaptive immune responses ^3^. However, many clinically significant pathogens have evolved strategies to resist or subvert complement-mediated attack, thereby enabling persistent colonization, immune evasion, and invasive disease^4,5^. Understanding how bacterial pathogens interact with and modulate complement activity is essential to uncovering mechanisms of immune resistance and to guiding the development of more effective vaccines and therapeutics. Two major human pathogens of particular concern in this context are *Streptococcus pneumoniae* and *Streptococcus pyogenes*, both of which cause significant global morbidity and mortality despite existing public health interventions ^6,7^. Much of our understanding of streptococcal pathogenesis and immune evasion has been derived from murine models, which have provided important insights into the mechanisms underlying infection. However, differences between human and murine immune systems may limit the extent to which these models recapitulate molecular host-pathogen interactions occurring during human infections ^8–10^.

*S. pneumoniae* a is Gram-positive strictly human pathogen. In 2019 alone, it was estimated to have caused over 800 000 deaths^11^. Commonly carried asymptomatically in the upper respiratory tract, particularly in children, *S. pneumoniae* can convert from commensal to pathogen, causing infections that range from sinusitis and otitis media to life-threatening pneumonia, sepsis, and meningitis ^7^. Its’ virulence repertoire includes the polysaccharide capsule, pneumolysin and DNases such as EndA ^12,13^. Conjugate vaccines, such as the 23-valent polysaccharide vaccine (PPSV23), 13-valent conjugate vaccine (PVC13), 15-valent (PCV15) and 20-valent (PCV20) conjugate vaccine have been developed against *S. pneumoniae* and lowered disease incidence ^7^, but challenges persist due to limited serotype coverage, and with over 98 known serotypes, expanding vaccine coverage remains both logistically difficult and economically expensive ^14,15^.

*S. pyogenes,* also called Group A Streptococcus (GAS), is a Gram-positive, strictly human pathogen capable of causing a broad clinical spectrum ranging from self-resolving and mostly mild infections such as pharyngitis and impetigo, to severe invasive infections such as necrotizing fasciitis, streptococcal toxic shock syndrome, and sepsis ^6^. Untreated infections can lead to immune-mediated sequelae including acute rheumatic fever, rheumatic heart disease, and glomerulonephritis ^16,17^. A 2005 global burden study estimated GAS to be responsible for 727 million non-invasive and 1.78 million severe cases annually, resulting in over 500,000 deaths, including 233,000 from rheumatic heart disease ^9,10^. Its pathogenicity is orchestrated by an array of virulence factors: surface proteins such as the family of M-protein and protein H and secreted proteins ^16^ including the streptococcal inhibitor of complement (SIC) ^18–20^, the *Streptococcus pyogenes* nuclease A (SpnA) ^21^, the complement evasion factor (CEF) ^22^, the pore-forming toxin streptolysin (SLO) ^23,24^, the cysteine protease SpeB and the IgG-degrading enzyme of *S. pyogenes*, IdeS ^25,26^.

Despite its profound public health impact and extensive study, no licensed vaccine is currently available for *S. pyogenes*. Developing a broadly effective vaccine against *S. pyogenes* has remained challenging, due to the risk of autoimmune complications and the difficulty of assessing protective immunity in a strictly human-adapted pathogen ^6^. A majority of clinical candidates target the surface-associated M-proteins, including the 30-valent N-terminal peptide vaccine from StreptAnova ^23^ or the MJ8VAX which targets the J8 epitope ^27^. Several non M-protein vaccine formulas have also emerged. These candidates are based on conserved antigens such as SLO, *S. pyogenes* Adhesion and Division Protein (SpyAD), streptococcal cysteine protease A (ScpA) and chemokine protease (SpyCEP) which have been formulated as multicomponent vaccines to broaden strain coverage^6,21,28,29^. While these candidates have shown promising results in preclinical trials, their protective efficacy needs to be further investigated as it is still unknown which virulence factors, secreted or cell surface-bound, lead to a protective immunity.

One of the conserved virulence strategies across both species is the secretion of DNases. These proteins target neutrophil extracellular traps (NETs), preventing bacterial entrapment and enhancing bacterial dissemination. In *S. pyogenes*, the prophage-encoded Sda1 is a highly active DNase associated with invasive M1T1 strains ^30,31^, enhancing resistance to neutrophil-mediated killing and promoting systemic dissemination. SpnA further contributes to NET degradation and streptococcal virulence by binding the human C5b67 assembly intermediate, sterically hindering the assembly of a functional MAC ^21^. Similarly, *S. pneumoniae* expresses EndA, a membrane-anchored nuclease with dual functionality: while essential for natural genetic transformation, EndA also degrades NETs during infection, contributing to pneumococcal virulence ^32,33^. The deployment of DNases by both species reflects a conserved strategy to neutralize extracellular immune traps and promote persistence within the host.

Here, we investigate whether the streptococcal DNases EndA and Sda1 possess virulence functions beyond NET degradation, focusing on their potential interactions with host blood plasma proteins. Using affinity-purification mass spectrometry (AP-MS) we define and compare the interactomes of Sda1 and EndA and benchmark them against SpnA. To our surprise, all three nucleases demonstrate moonlighting functions as complement binders and possible inhibitors. To place our findings in a functional context, we performed enzyme-linked immunosorbent assays (ELISAs) to gain orthogonal confirmation of the host complement proteins targeted, and to determine which complement pathway is inhibited by these nucleases. We further performed hemolysis inhibition assays to determine if Sda1 and EndA sterically obstructs MAC assembly, hence inhibiting the proper function of the human complement system. This integrated approach allowed us to assess not only whether these nucleases interfere with complement activity but also identify specific factors targeted by each enzyme, broadening our understanding of DNase-mediated virulence functions in *S. pyogenes* and *S. pneumoniae*.

## MATERIALS AND METHODS

### Proteins

The cloning, expression and purification of recombinantly expressed SpnA and superfolder green fluorescent protein (sfGFP) has been described before^34,35^. The proteins EndA and Sda1 were expressed as inactive mutants, EndA with a mutation at H160G and Sda1 at H188G. A 6xHiS-HA-StrepII-TEV (histidine-hemagglutinin-StrepII-Tobacco Etch Mosaic Virus protease recognition site) tag on the C-terminus was added to EndA (H160G) (UniProt ID: P0A3S3) and Sda1(H188G) (UniProt ID: Q675N6) ^33,36^. These were ordered as a synthetic construct from Genscript and cloned into the pET26b(+) vector. EndA and Sda1 were expressed in LB (Liquid Broth) in *E. coli* TURNER (DE3) cells. Expression induced with 1 mM IPTG at OD_600_= 0.8 and expressed for 18 hours. The cells were harvested after 18 h and resuspended in wash buffer (25 mM Tris-HCl, 100 mM NaCl, pH 8.0) supplemented by Complement Protease inhibitor EDTA-free tablets (Roche) as well as DNase I and were then passed two times through a French Pressure Cell at 18,000 psi. The lysate was cleared by ultracentrifugation (Ti 50.2 rotor, 60 min, 4°C, 45,000 rpm). Subsequently, the soluble lysate was syringe-filtered with a 0.45 µm for Sda1, respectively a 0.25 µm filter for EndA and loaded a HisTrap HP column (Cytiva). A 5 mL HisTrap HP column (Cytiva) was connected to an ÄKTA Pure chromatography system (Cytiva). The column was washed with H_2_O and then 5 column volumes of phosphate buffer (50 mM sodium phosphate, 300 mM NaCl, 20 mM imidazole, pH 8.0) followed by equilibration by imidazole phosphate buffer (50 mM sodium phosphate, 300 mM NaCl, 500 mM imidazole, pH 8.0). The sample was applied and the column was washed with 20 column volumes with the phosphate buffer. The bound protein was eluted with a 0-100% gradient with the imidazole phosphate buffer at a flow rate of 2.5mL/min. Fractions containing the proteins were pooled and the purified EndA and Sda1 was mixed with TEV protease at a 1:20 enzyme:substrate ratio and DTT was added to a final concentration of 1 mM and the digestion was incubated overnight at 16°C. The TEV-cleaved proteins were purified on a HisTrapHP column with the flowthrough and wash fractions were collected. Subsequently, affinity-purified and protease-cleaved EndA and Sda1 were further purified by size-exclusion chromatography using HiLoad 26/600 Superdex 75 pg column (GE Healthcare) in 25 mM Tris HCl, 100 mM NaCl pH 8. All proteins were expressed and purified at the Lund Protein Production Platform (LP3), Lund, Sweden.

For the enzyme-linked immunosorbent assays (ELISAs), the purified complement system proteins C5 (A120), C6 (A123), C5b6 (A122), C7 (A124), C8 (A125), C9(A126) and the sC5b-9 (A127) complex were purchased from Complement Technology, Inc. (Texas, USA).

### Plasma, serum and whole blood

Pooled human plasma from healthy donors and pooled BALB/c mouse plasma was purchased from Innovative Research (MI, USA). Single donor human serum and whole blood were collected from healthy volunteers. The blood was centrifuged at 3000 × g for 10 min to collect the erythrocytes. Ethical approval was obtained from the local ethics committee (approval 2025-05426-01), and the study was conducted according to the Declaration of Helsinki.

### Affinity-purification (AP-MS)

The affinity-purification experiments have been described ^21^. Strep-Tactin beads (IBA Life Sciences) were equilibrated in PBS-buffer (1x PBS pH 7.4, 1 mM CaCl_2_, 5 mM MgSO_4_) and charged with 100 μg each of recombinant, affinity-tagged EndA, Sda1 and sfGFP as five replicates. Pooled normal human plasma (100 μL) or BALB/c mouse plasma (100 μL) was incubated with the protein-charged beads at 30 min, 37°C, 500 rpm. The beads were washed with 10 mL ice-cold PBS-buffer as above, and proteins eluted with 180 µL of 5 mM biotin in PBS-buffer. The samples were precipitated with a final concentration of 25% trichloroacetic acid (TCA), washed with acetone and dried in a speedvac prior to sample preparation for mass spectrometry.

### Sample preparation for mass spectrometry (MS)

The affinity-purified samples were resuspended in 8 M urea-100 mM ammonium bicarbonate (ABC), cysteine-bonds reduced using 5 mM tris(2-carboxyethyl) phosphine hydrochloride (TCEP) (60 min, 37°C, 800 rpm) and alkylated with 5 mM iodoacetamide (30 min, 22 °C). The samples were subsequently diluted with ABC to a UREA concentration of below 1.5 M. Sequencing grade trypsin (Promega) was added to the samples for digestion (18 h, 37°C, 1000 rpm). The samples were acidified with 10% formic acid (FA) to a final pH < 3.0 and purified using C18 reverse phase spin columns according to manufacturer’s instructions (The Nest Group, Inc.). The purified peptides were dried in a speedvac until completely dry. Followed by sample cleanup with SP3-beads (Sera-Mag beads Thermo Scientific) ^37^.

### Liquid chromatography tandem mass spectrometry (LC–MS/MS)

LC-MS/MS in DDA (data dependent acquisition) mode was performed on an Exploris™ 480 hybrid quadrupole-Orbitrap mass spectrometer (Thermo Fisher) system connected to a NeoVanquish pump (Thermo Fisher Scientific). The peptides were trapped on an Acclaim™ PepMap™ 100 C18 HPLC pre-column (75 µm × 2 cm, with 3 µm particle size; Thermo Fisher) and separated on an Easy-Spray PepMap™ RSLC C18 analytical column (75μm x 25 cm, with 2 µm particle size; Thermo Fisher). Mobile phases consisted of 0.1% FA in H_2_O (A) and 80% acetonitrile/0.1% FA (B). Peptides were separated at a flow rate of 300 nL min^-1^ using a gradient of 5-25% B over 50 min, 25-32% B over 6 min and 32-45% B over 4 min. Subsequently followed by washing at 95% B for the remainder of the 70 min analysis.One full MS scan was acquired at a resolution of 120,000 of 375–1500 m/z, followed by MS/MS scans at a resolution of 15,000. Precursor ions were selected with a charge state of 2-6 and an isolation window of 1.3 m/z. The precursors were fragmented using higher-energy collisional dissociation (HCD) at a normalized collision energy of 30.

### Data analysis of affinity-purification experiments

The acquired spectra were analyzed against the reviewed human proteome (UniProt ID: UP000005640, containing 20,406 sequences) for the AP-MS samples in human plasma, and against the reviewed mouse proteome (UniProt ID: UP000000589, containing 17,224 sequences) for the AP-MS samples in mouse plasma. The sequence databases were supplemented with the sequences for sfGFP (UniProt ID: P42212), EndA (UniProt ID: P0A3S3), Sda1 (UniProt ID: Q675N6), and SpnA (UniProt ID: Q9A0J7), sequences for the affinity-tag (N-terminal or C-terminal), and were analyzed in MaxQuant 1.6.10.43. Full tryptic digestion was applied, allowing for up to 2 missed cleavages, with peptide length ranging from 6 to 144 amino acid residues. Carbamidomethylation of cysteine residues was defined as a static modification, while methionine oxidation was set as a variable modification. The precursor ions mass tolerance was set to 20 ppm for MS1 scans and 0.5 Da for MS2 scans. A false discovery rate (FDR) threshold of 1% was applied.

The proteomics data were filtered to include proteins identified by at least two unique peptides. Label-free quantification (LFQ) intensity values from MaxQuant were analyzed in Perseus (version 2.0.9.0). The data were Log2 transformed and median-normalized and filtered to retain proteins detected in at least three out of five replicates per bait protein. Missing values were imputed from a normal distribution (width 0.3, down shift 1.8). Differential abundance was assessed using a two-sample t-test with a 1% false discovery rate (FDR < 0.01) and S0 = 2 as thresholds for significance. Significant proteins were visualized using scatterplots in GraphPad Prism (version 10.4.1). Alternatively, for bar graphs, the data were Log2 transformed and bait-sum normalized. Normalized values were visualized as bar charts in GraphPad Prism, where individual replicate points and standard deviation overlaid to illustrate variability.

### Complement pathway inhibition assays

To qualitative determine the EndA, Sda1 and SpnA mediated fuctional inhibition of the classical and alternative complement pathways, we used the WIESLAB® Complement Screen, also as described before for CEF ^22^. For both pathways, 10 μM of each protein was mixed with 5 μl of single donor serum and incubated for 30 min at 37°C. For the classical pathway, the serum was diluted 1:101 in assay diluent and 100 µl was transferred to each well of a 96-well plate. For the alternative pathway, serum was diluted 1:18 and processed identically. Plates were incubated for 1 hour at 37°C and washed three times with 300 µL wash buffer. Subsequently, 100 µL conjugate was added and incubated for 30 minutes at room temperature. After another round of three washes, 300 µL each, 100 µL substrate solution was added and incubated in the dark for 30 min. Subsequently, absorbance was measured at 405 nm. Complement activity was calculated using the OD_405_ mean values for the sample, positive control (PC) and negative control (NC) using the kit formula and the data visualized in GraphPad Prism:

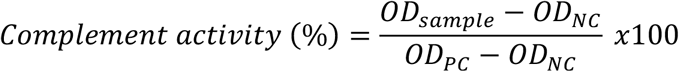

### Hemolytic assay

To test the effects of EndA and Sda1 as inhibitors of complement mediated cell lysis, EndA and Sda1 were diluted to 10 μM in 1x PBS, pH 7.4 buffer complemented with 1 mM CaCl_2_ and 5 mM MgSO_4_. The complement components C5b6, C7, C8 and C9 were all diluted separately in the reaction buffer; C5b6 to 20 μg/ml and the others to 10 μg/ml. For the hemolytic assay, all experiments were prepared as three technical replicates. Here, we used 50 μL of 5% human erythrocytes in reaction buffer, mixed with 25 μl of EndA, Sda1 or BSA and 25 μL of each complement component added sequentially, and followed by incubation for 30 min at 37°C. Erythrocytes mixed with water, with reaction buffer or with the complement components in reaction buffer only, were used as controls. The plates were centrifuged to pellet intact cells and cell membranes, and 100 µl supernatant from each well was transferred to a fresh plate to measure hemoglobin release as a measure of cell lysis. The percentage of lysis was calculated according to Fernie-King *et al.* ^19^. Previous hemolysis data of SpnA and BSA was reanalyzed from A. Bennig *et al*. for comparison ^21^.

### Enzyme-linked immunosorbent assay (ELISA)

To investigate binding of EndA and Sda1 to individual components of the complement system MAC, triplicate wells were coated with purified C5, C6, C5b6, C7, C8, C9 or sC5b-9 at 2 µg/ml as above or buffer only as a negative control and incubated over night at 8°C. Wells were blocked using 1x PBS, pH 7.4 complemented with 0.05% Tween-20 (PBS Tween) containing 2% BSA for 60 min, room temperature (RT) before affinity-tagged EndA and Sda1 at 10 µg/ml were added and incubated for 2 h, RT. Following washing with PBS-T binding of EndA or Sda1 complement proteins was detected with 6x-His Tag Polyclonal Antibody, HRP (Invitrogen) diluted 1:5000 in blocking buffer for 60 mins, RT. Wells were washed with PBS-T before the addition of 3,3’,5,5’-Tetramethylbenzidine (TMB) substate. The reaction was allowed to proceed for 15 min, RT, in the dark before reading absorbance at 450 nm.

## RESULTS

### Host-plasma interaction networks reveal human-specific interactions with complement

To define the plasma interactomes of the streptococcal virulence factors EndA and Sda1, we performed AP-MS with pooled normal human or mouse blood plasma. Here, recombinant EndA and Sda1 were produced with a C-terminal affinity tag and used as bait proteins. The sfGFP were used as negative control. For EndA we were able to identify 16 high confidence interactions in human plasma, respectively 15 for Sda1 when filtered against sfGFP at an FDR < 0.01 and a Log2-fold enrichment of 2 (**Figures 1A-B**). The significantly enriched proteins could be grouped into two main categories of complement system associated proteins, including complement inhibitors, as well as other plasma proteins. Interestingly, most of the complement proteins targeted belong to the MAC assembly. This pattern mirrors the interactome we have previously observed for SpnA ^21^, and suggested a novel, conserved function for streptococcal nucleases in complement binding and possible complement inhibition. Direct comparison of the EndA and Sda1 interactomes revealed both common and nuclease-specific binding profiles (**Figure 1C**). Whereas EndA binds the complement system-associated clusterin (CLUS) and mannan-binding lectin serine protease 1 (MASP1) at an apparent higher affinity than Sda1, Sda1 interacts specifically to several human plasma proteins, including the immunoglobulin G subclasses IgG1 (IGHG1) and IgG4 (IGHG4) (**Figure 1C**). To place these interactions in context, we compared the interactomes of EndA and Sda1 generated here to that of SpnA described previously ^21^. This analysis revealed that SpnA has a much richer human plasma interactome as compared to EndA and Sda1, with 24 unique plasma proteins targeted in contrast to 7 or 8, respectively (**Figure 1D**). This difference in the human plasma interactomes might in part be explained by the size difference of the proteins with SpnA close to 100 kDa, and EndA and Sda1 at 27 kDa and 40 kDa, respectively; allowing for more interaction interfaces at the SpnA surface. Regardless, the analysis highlight both the differences and similarities between the host proteins targeted by these three nucleases, such as the interactions to the complement inhibitors vitronectin (VTCN) and CLUS being unique to EndA and SpnA, while hornerin (HORN) being specific to Sda1 and SpnA (**Figure 1D**). To our surprise, the only proteins targeted by all three nucleases were those belonging to the MAC assembly: C5, C6, C7, C8A, C8B, C8G and C9. Together, these comparisons underscore that while all three nucleases converge on the MAC, each factor also engages distinct subsets of host proteins.

**Figure 1.**
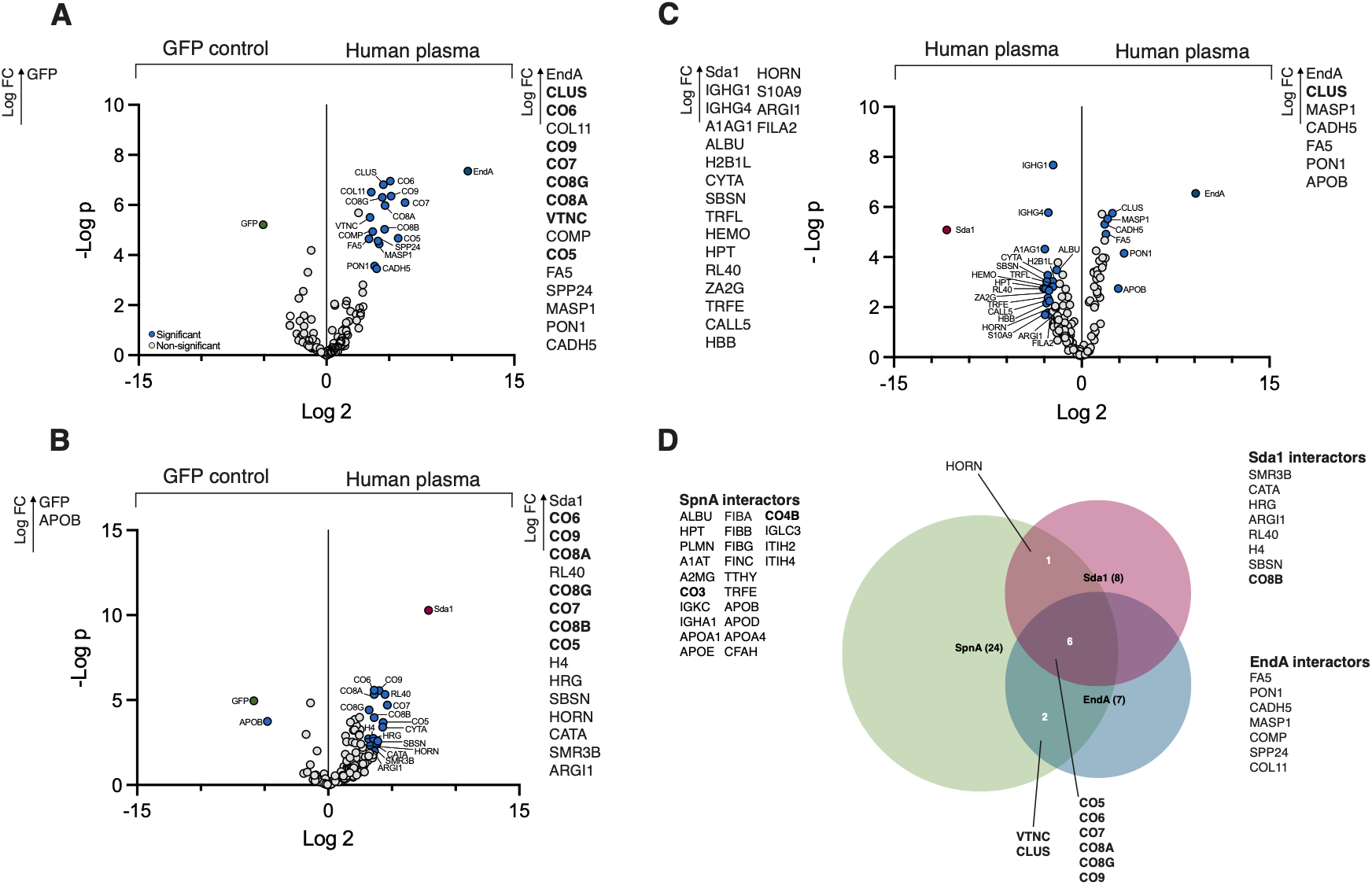
Human plasma interactomes of EndA and Sda1. **A)** Volcano plot analysis comparing the human plasma interactome of EndA to sfGFP; **B)** Sda1 to sfGFP and **C)** Sda1 to EndA when filtered against sfGFP at an FDR < 0.01 and a Log2-fold enrichment of 2. Significantly enriched proteins are indicated on both sides of the plot in Log2 fold-change (FC) order. Key human interactors of specific interest are marked in bold. **D)** Venn diagram comparing significant proteins in the EndA, Sda1 and SpnA interactomes. The SpnA data is reanalyzed from A. Bennig *et al* ^21^.

### The mouse plasma interactomes differ sharply from the human interactomes

To assess nuclease species specificity, we compared our human AP-MS data to an AP-MS interactome using mouse blood plasma. In striking contrast to the human interactomes, we could not identify any significant EndA interactors when filtered against sfGFP at an FDR <

0.01 and a Log2-fold enrichment of 2 (**Figure 4A**), indicating a strictly human specific function. For Sda1 we were able to identify 8 significant interactors when filtered against sfGFP, where APOA1 was the most abundant interactor (**Figure 4B**) followed by plasminogen (PLMN), APOE, the immunoglobulin G subclass IgG3 (IGHG3), CLUS and the mannose-binding protein C (MBL2). A comparison of the two nucleases EndA and Sda1 filtering at an FDR < 0.01 and a Log2-fold enrichment of 2, further demonstrated the lack of EndA-specific interactors in mouse plasma (**Figure 4C**).

None of the identified mouse interactors belonged to the complement system MAC, underscoring that the complement targeting by these nucleases is specific to the human species. Consistent with this, the overall binding profiles were sparse compared to the human interactomes. These findings highlight a pronounced species dependence: whereas human plasma interactomes were enriched for multiple high-confidence targets, those in mouse plasma were largely lacking (**Figure 1 and 4**). Extending the analysis to all three nucleases further revealed minimal overlap between the interactomes of SpnA, EndA and Sda1 in mouse blood plasma (**Figure 4D**). No protein was shared across all three nucleases, and only a few were shared pairwise: APOE, APOA1 and CLUS between Sda1 and SpnA, and serotransferrin (TRFE) between EndA and SpnA (**Figure 4D**). Interestingly, TRFE has for some species, including streptococci, been demonstrated to serve as an iron source, where the microbe or parasite captures the TRFE and extract its iron for their own use ^38–42^. This highlights a pronounced species dependence and has important implications for interpreting murine models of infection ^8,9^.

### Quantitative comparison of host protein enrichment by nucleases

To validate and quantify enrichment of selected human plasma proteins identified in AP-MS, we compared key proteins quantified across EndA, Sda1 and the SpnA data from A. Bennig *et al.* was reanalyzed^21^. For each protein of interest, the LFQ intensities were Log2 transformed and normalized to the bait by subtracting the corresponding bait from each replicate (**Figure 2**, **Figure 3**). This analysis confirmed nuclease-specific trends observed in the volcano plots (**Figures 1 and 4**). EndA and SpnA pulldown C1R, C1S, C1QC, C4B, C4BPA and MASP1 more, while SpnA pulldown alone C4A, compared to both EndA and Sda1 (**Figure 1**). Interestingly EndA binds IGHG1 and IGHG3 but not IGHG2 or IGHG4 (**Figure 2**), furthermore, our data shows that EndA binds C1r, C1s, C1qB and C1qC as opposed to Sda1 (**Figure 3**) which suggests that EndA acts primarily by interfering with initiation of the classical complement pathway. Furthermore, the three nucleases EndA, Sda1 and SpnA exhibit a similar level of enrichment of MAC components. We also observed a comparable enrichment of VTNC and CLUS, both of which are established regulators of complement system (**Figure 3**) and part of the soluble MAC (sC5b-9 complex) ^43,44^.

**Figure 2.**
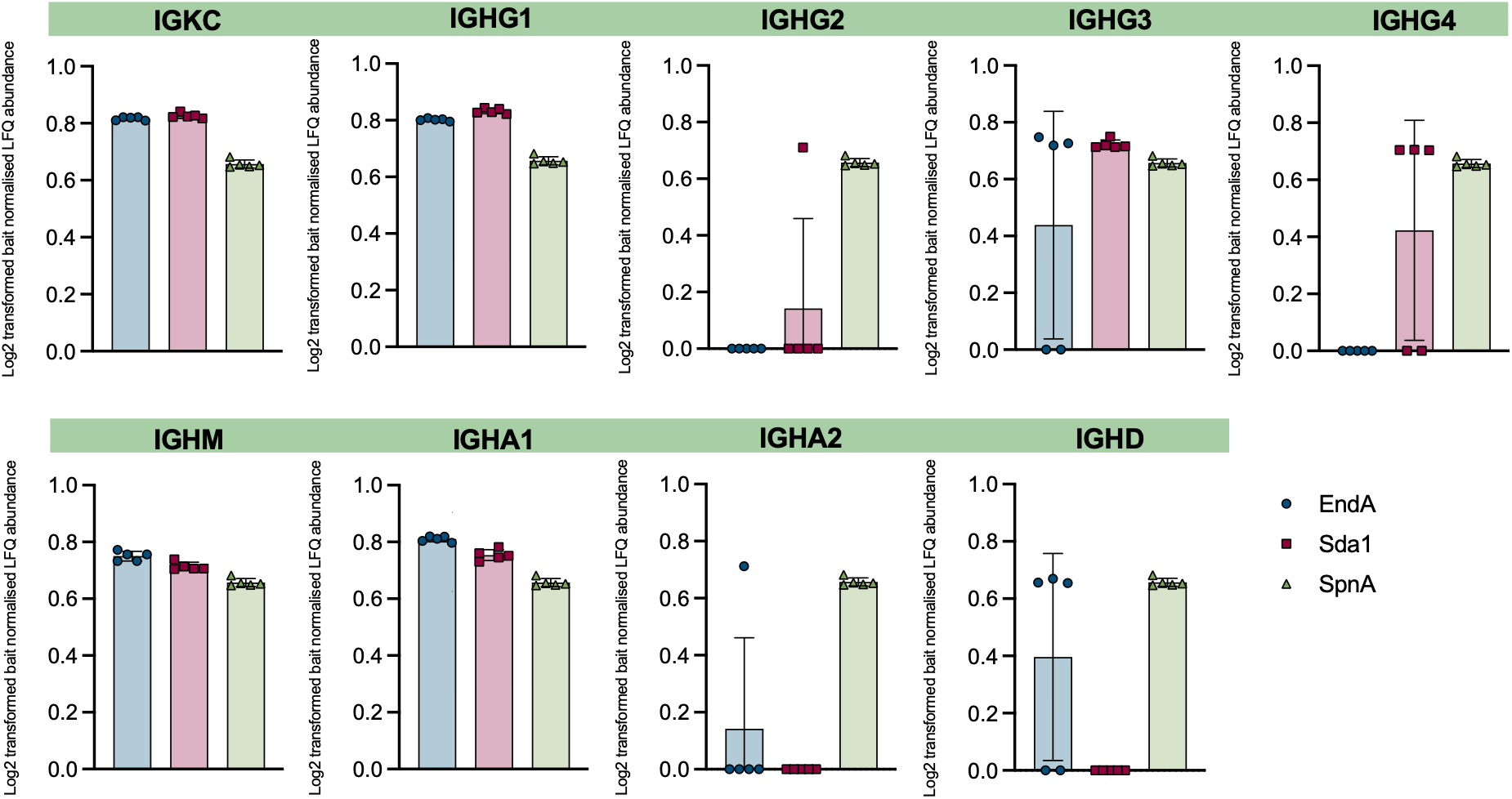
Quantitative comparison of all immunoglobulins enriched by EndA, Sda1 and SpnA. Bar charts showing Log2-transformed bait normalized LFQ abundances of selected proteins across five independent replicates. Error bars represent standard deviation across replicates. The SpnA data is reanalyzed from A. Bennig *et al* ^21^.

**Figure 3.**
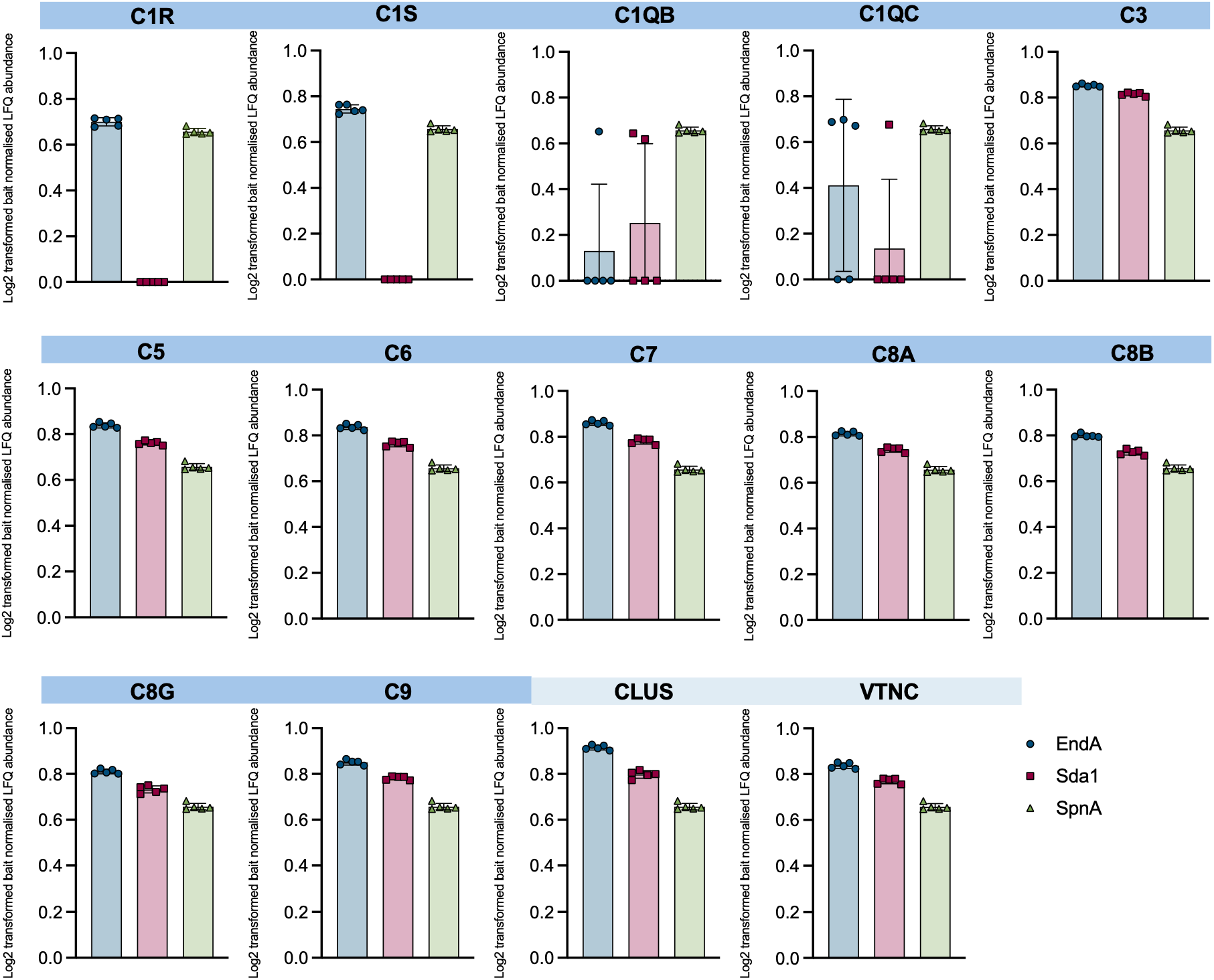
Quantitative comparison of selected human plasma proteins enriched by EndA, Sda1 and SpnA. Bar charts showing Log2-transformed bait normalized LFQ abundances of selected proteins across five independent replicates. Error bars represent standard deviation across replicates. The SpnA data is reanalyzed from A. Bennig *et al* ^21^.

**Figure 4.**
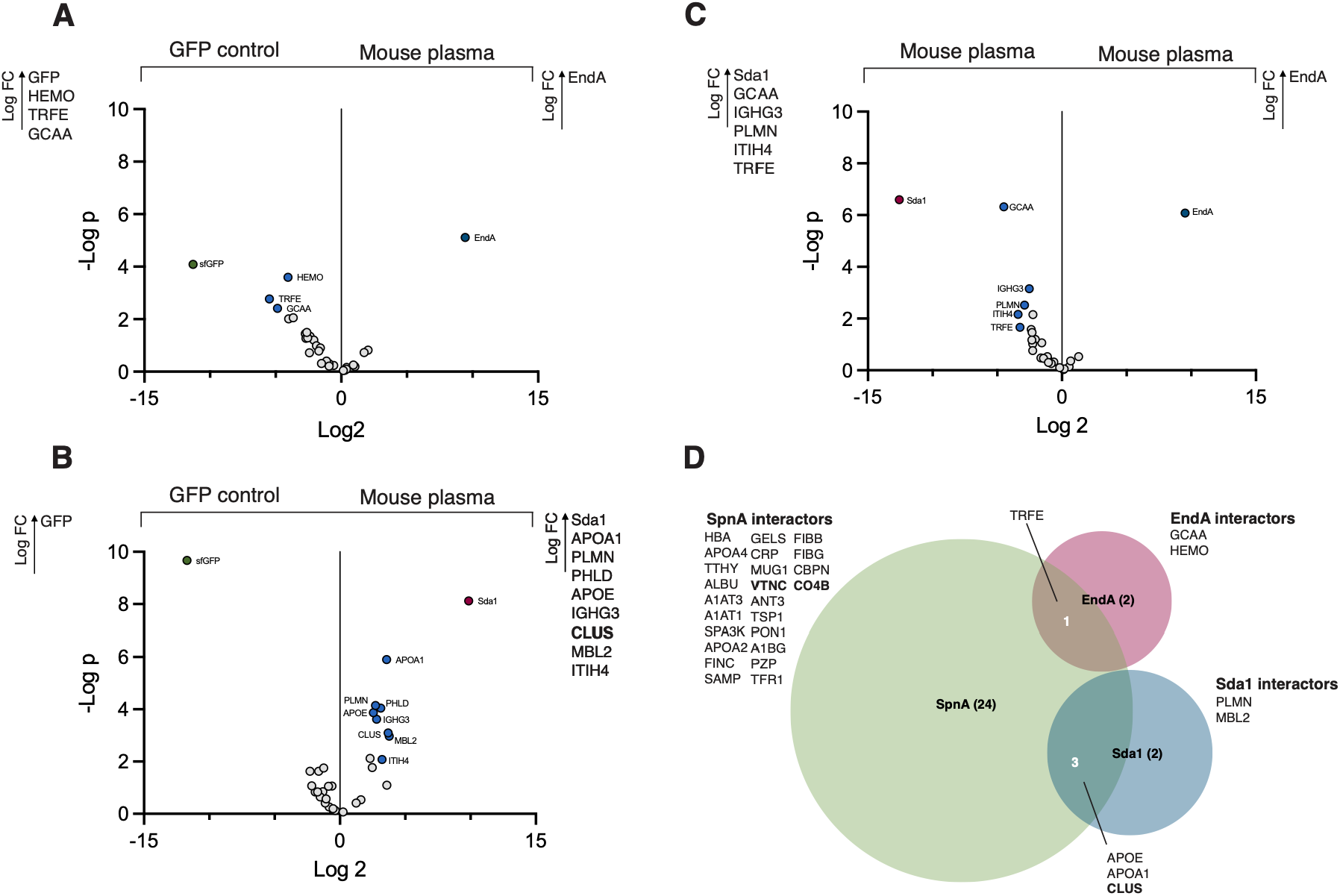
Mouse plasma interactomes of EndA and Sda1. **A)** Volcano plot analysis comparing the mouse plasma interactome of EndA to sfGFP; **B)** Sda1 to sfGFP and **C)** Sda1 to EndA when filtered against sfGFP at an FDR < 0.01 and a Log2-fold enrichment of 2. Significantly enriched proteins are indicated on both sides of the plot in Log2 fold-change (FC) order. Key mouse interactors of specific interest are marked in bold. **D)** Venn diagram comparing significant proteins in the EndA, Sda1 and SpnA interactomes. Minimal overlap is observed, with only a few pairwise shared proteins. The SpnA data is reanalyzed from A. Bennig *et al* ^21^.

### EndA and Sda1 bind to individual complement proteins

Previous studies have shown that streptococcal proteins, SIC and SpnA disrupt terminal complement complex formation by blocking insertion of the C5b67 complexes ^19,21^, whereas CEF has also been show to inhibit the complement medicated cell lysis ^22^. To validate the interactions identified by AP-MS and investigate whether EndA and Sda1 bind specific complement proteins, individual complement components, C5, C5bC6, C6, C7, C8, C9 and sC5b-9 complex that were immobilized on ELISA plates and incubated with affinity-tagged Sda1 or EndA. Sda1 produced elevated absorbance signals for C5, C5bC6, C8, C9 and sMAC relative to the negative control, whereas the signal observed for C7 was comparable to background levels (**Figure 5A**). Similarly, EndA generated elevated signals for multiple complement proteins, suggesting interaction with several components of the terminal complement pathway (**Figure 5B**). Together, these findings are consistent with interactions between EndA and Sda1 and several components of the terminal complement pathway.

**Figure 5:**
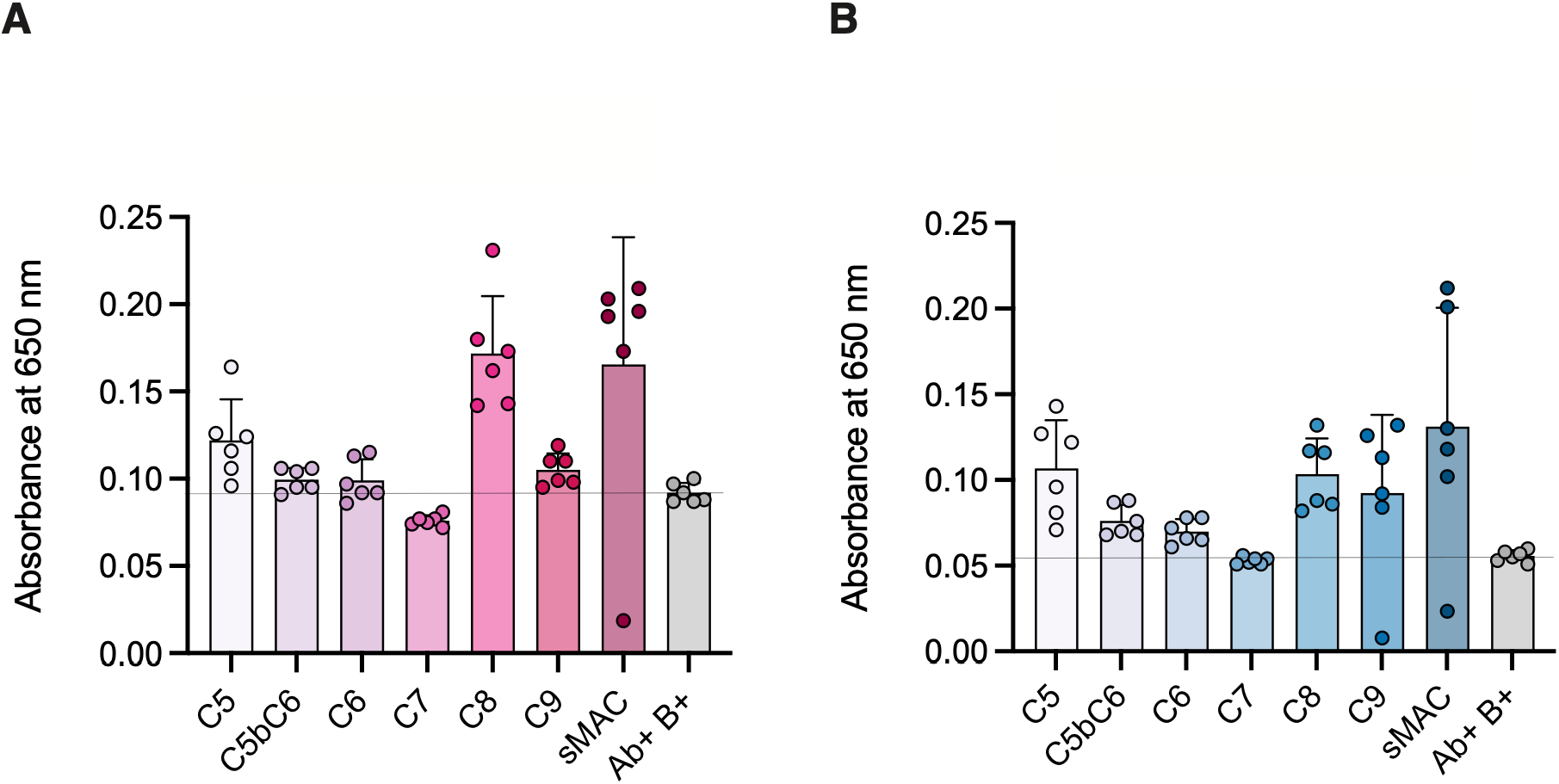
ELISA of Sda1 and EndA together with individual human complement proteins. Plates were coated with individual human complement proteins or the sMAC and incubated with affinity-tagged Sda1 or EndA. Binding was detected using an anti-6xHis HRP-conjugated antibody and binding was quantified by measuring absorbance at 650 nm. **A)** Binding of Sda1 with the individual complement proteins or sMAC. **B)** Binding of EndA with the individual complement proteins or sMAC. Wells containing antibody and bait protein in PBS supplemented with 1mM CaCl_2_ and 5mM MgSO_4_ pH 7.4, served as negative control. Data is presented as mean ± SD of triplicate wells.

### EndA inhibits the formation of the MAC and prevents hemolysis while Sda1 does not

To investigate whether the observed interactions translated into functional inhibition of complement activity, a complement-mediated hemolysis assay was performed using a modified version of the method described by Fernie-King et al ^19,21^. Human erythrocytes were incubated with complement in the presence of EndA, Sda1, SpnA (positive control) or BSA (negative control). Treatment with EndA reduced hemolysis to approximately 30 %, compared to 44% in Sda1, 20% in SpnA and 46% in the BSA treated control (**Figure 6A**). These results indicate that EndA and SpnA inhibits MAC-mediated lysis of erythrocytes, while Sda1 exhibits little to no inhibitory activity under the conditions tested ^21^.

**Figure 6.**
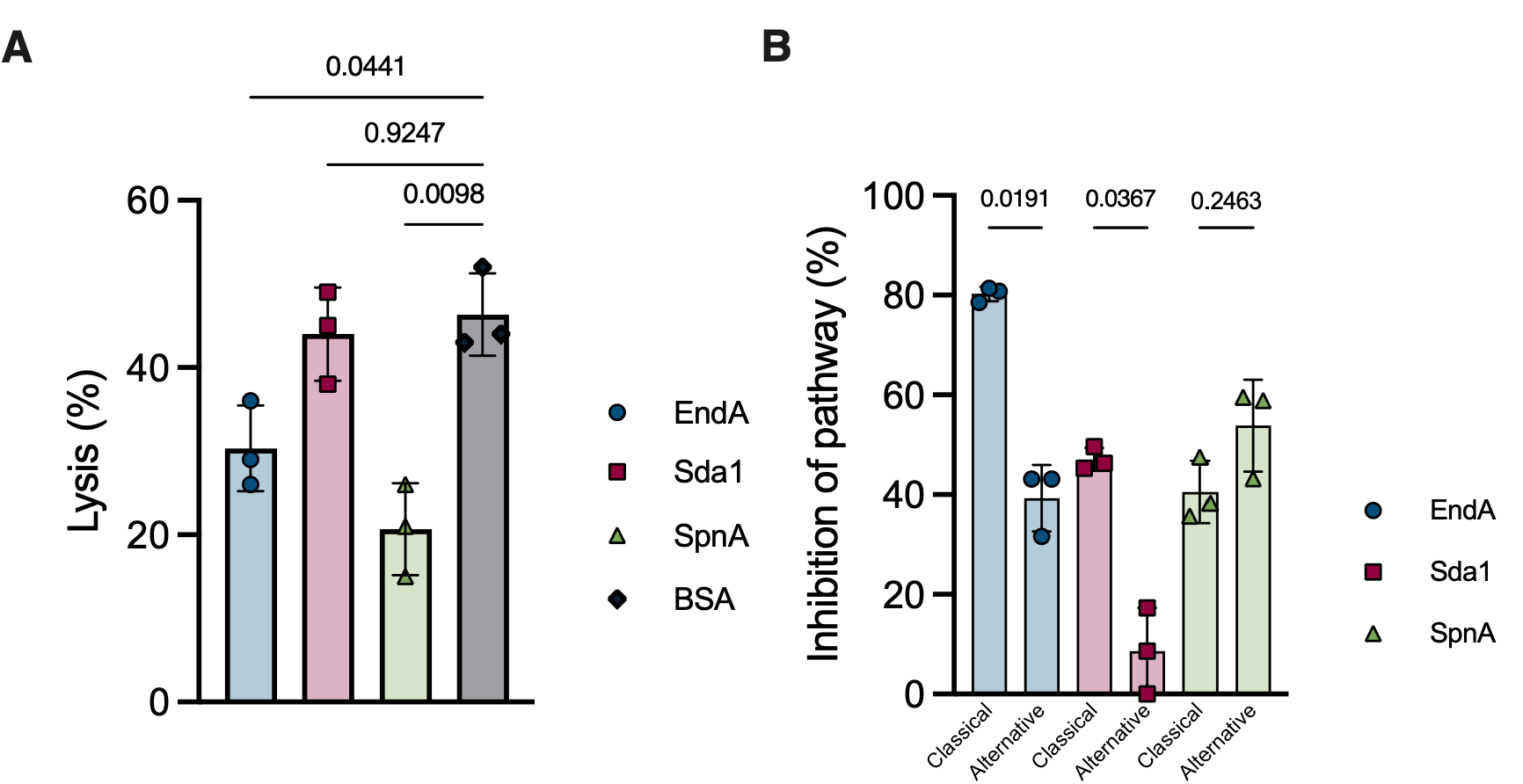
Hemolysis and complement assay EndA, Sda1, SpnA and BSA. **A)** Three individual samples of each protein were measured for lysis of human erythrocytes. The assay indicates that lysis was inhibited in the presence EndA and SpnA. Sda1 and BSA display no protection in lysis. A one-way ANOVA was used for statistical analysis. Statistical significance is indicated as follows: *P* > 0.05. **B)** Complement activity was assessed in the presence of EndA, Sda1 and SpnA using the (kit). Activation of the classical pathway and alternative pathway was quantified measuring absorbance at 405 nm. Bars represent the mean of triplicate measurements, and error bars indicate standard deviation. A one-way ANOVA was used for statistical analysis. Statistical significance is indicated as follows: *P* > 0.05. The SpnA data is reanalyzed from A. Bennig *et al* ^21^.

### EndA and Sda1 inhibit the classical pathway more than the alternative

Based on our results, we wanted to determine what complement pathway is targeted by EndA, Sda1 or SpnA. To do this we performed the WIESLAB® Complement Screen for the classical and alternative pathways, as EndA (PDB: 3OWV) and Sda1 (PDB: 5FGW) are proteins lacking carbohydrate moieties, the lectin pathway was not investigated. To benchmark the inhibitory effect of EndA and Sda1, we compare them to an already established inhibitor of the complement, SpnA. Both EndA and Sda1 inhibited the classical pathway more effectively than the alternative pathway (**Figure 6B**). EndA reduced classical pathway activity by approximately 80% compared with 40% inhibition for the alternative pathway. Similarly, Sda1 reduced the classical pathway activity by approximately 45%, while reducing alternative pathway activity by only 10%. In contrast, SpnA displayed a different inhibition profile, reducing alternative pathway activity by approximately 55% and classical pathway activity by 40%. The difference in inhibition between the classical and alternative pathways was statistically significant for both EndA and Sda1 (*P < 0.05*) indicated a preferential effect on the classical pathway as opposed of SpnA (**Figure 6B**).

## DISCUSSION

Gram-positive bacteria, including *S. pyogenes* and *S. pneumoniae*, continue to remain big health burdens, responsible for a wide spread of non-invasive and invasive diseases. Their persistence and high infection numbers are closely linked to their large virulence factor arsenal and ability to evade the human immune system, as demonstrated here. While Gram-positive bacteria are thought to resist complement-mediated lysis due to their thick peptidoglycan layers, recent evidence shows that MAC can assemble on their surfaces and under specific conditions and adopt pore-forming conformations ^1,2^.

Despite the availability of antibiotics and conjugate vaccines, both pathogens continue to pose significant challenges in clinical settings due to the rise of antimicrobial resistance and limited stereotype coverage for existing vaccines. This highlights the importance and need for alternative or complementary therapeutic strategies. Virulence factors that are secreted or surface-bound are attractive targets for therapeutic intervention, as these often play a crucial role in pathogen survival, immune evasion and host colonization, making them interesting targets for therapeutic intervention. Our findings contribute to this growing field of research by further characterizing the functions of EndA and Sda1.

In this study, we present a comprehensive host-pathogen interactome of EndA and Sda1 by integrating proteomic mapping with functional assays to uncover novel interactions and functions. EndA and Sda1 target the human MAC, a finding only observed in the human interactome as the nucleases do not show any binding to mouse complement proteins. This suggests that interaction with the complement proteins is conserved in humans and reflects a host-specific evolutionary pressure. Animal models, particularly murine models, have contributed substantially to our understanding of streptococcal pathogenesis and host immunity. However, species-specific differences between the two immune systems, can limit the extent to which these models capture and represent interactions during human infections. This is particularly relevant in studies investigating host-pathogen interactions, with focus on differences in abundance, sequence or function of host proteins that may influence the molecular interactions between host and pathogen. Thus, the characterization of the human host-pathogen interactome is relevant and central in developing future therapeutic targets.

Functional validation reveals that EndA from *S. pneumoniae* is a new complement system inhibitor, following the functional pattern of SpnA, and suggested to be active in the prevention the insertion of the MAC into cell membranes. In contrast, Sda1 does not exhibit the same inhibitory activity despite sharing structural similarities and host-pathogen interactomes with EndA. Interestingly, SpnA, EndA and Sda1 all interact with the C1QB and C1QC subunit of the C1 complex, suggesting a potentially shared mechanism of interference with the classical complement pathway activation. However, differences were observed in their interaction profiles, as Sda1 did not interact with the C1R or C1S proteases. Despite these differences, all three nucleases demonstrate activity inhibition on the classical pathway. This suggests that direct interaction with C1R or C1S might be necessary for classical pathway inhibition. Instead, interaction with one of the C1Q subunits might be sufficient ^45^. This functional difference can be a result of substrate specificity or additional co-factor interactions that were not captured in our current assays.

In conclusion, this study provides a comprehensive characterization of host-pathogen interactions involving *S. pneumoniae* and *S. pyogenes*, further expanding our understanding of the molecular mechanisms in streptococcal infection and immune evasion. These findings highlight the complexity of the host-pathogen interplay and the importance of investigating such interactions in a human context. By defining these interactions, this work establishes a resource for future mechanistic studies that may contribute to novel therapeutics and vaccine targets.

## ACKNOWLEDGEMENTS

Support to LJH from Paul and Caroline Sverdrup is gratefully acknowledged. This work was supported by grants from the Swedish Research Council (2022-03860), the Royal Physiographic Society in Lund, Stiftelsen Clas Groschinskys Foundation, and the Foundations of Crafoord and Alfred Österlund to LJH. We acknowledge Protein Production Sweden (PPS) for providing facilities and experimental support. PPS is funded by the Swedish Research Council as a national research infrastructure. Support from the Swedish National Infrastructure for Biological Mass Spectrometry (BioMS) is gratefully acknowledged. The authors thank Lars Björck for valuable discussions and critical comments on the manuscript.

## AUTHOR CONTRIBUTIONS

I.A.B. and L.J.H. conceptualized the manuscript. The methodology was jointly designed by I.A.B, S.K. and L.J.H. I.A.B, S.K. and L.J.H. conducted the laboratory experiments and data collection. Data analysis and interpretation were carried out by I.A.B., S.K. and L.J.H. I.A.B., S.K. and L.J.H. made the figures. L.J.H. supervised the work. I.A.B wrote the initial draft of the manuscript. All authors have reviewed and approved the final version of the manuscript.

## DATA AVAILABILITY STATEMENT

The affinity-pulldown data have been deposited to the ProteomeXchange consortium via the MassIVE partner repository https://massive.ucsd.edu/ with the dataset identifier MSV000104147. The plasmids generated in this study have been deposited into the Addgene repository with the following IDs: 263752 (EndA) and 263753 (Sda1).

## REFERENCES

1. Berends, E. T. M., Dekkers, J. F., Nijland, R., Kuipers, A., Soppe, J. A., Van Strijp, J. A. G., & Rooijakkers, S. H. M. (2013). Distinct localization of the complement C5b-9 complex on Gram-positive bacteria: Localized C5b-9 deposition on Gram-positive bacteria. Cellular Microbiology, 15(12), 1955–1968. 10.1111/cmi.12170

2. Khakzad, H., Happonen, L., Tran Van Nhieu, G., Malmström, J., & Malmström, L. (2021). In vivo Cross-Linking MS of the Complement System MAC Assembled on Live Gram-Positive Bacteria. Frontiers in Genetics, 11, 612475. 10.3389/fgene.2020.612475

3. Dunkelberger, J. R., & Song, W.-C. (2010). Complement and its role in innate and adaptive immune responses. Cell Research, 20(1), 34–50. 10.1038/cr.2009.139

4. Barber, M. F., & Fitzgerald, J. R. (2024). Mechanisms of host adaptation by bacterial pathogens. FEMS Microbiology Reviews, 48(4), fuae019. 10.1093/femsre/fuae019

5. Su, M. S.-W., Cheng, Y.-L., Lin, Y.-S., & Wu, J.-J. (2024). Interplay between group A *Streptococcus* and host innate immune responses. Microbiology and Molecular Biology Reviews, 88(1), e00052–22. 10.1128/mmbr.00052-22

6. Brouwer, S., Rivera-Hernandez, T., Curren, B. F., Harbison-Price, N., De Oliveira, D. M. P., Jespersen, M. G., Davies, M. R., & Walker, M. J. (2023). Pathogenesis, epidemiology and control of Group A Streptococcus infection. Nature Reviews Microbiology, 21(7), 431–447. 10.1038/s41579-023-00865-7

7. Narciso, A. R., Dookie, R., Nannapaneni, P., Normark, S., & Henriques-Normark, B. (2025). Streptococcus pneumoniae epidemiology, pathogenesis and control. Nature Reviews Microbiology, 23(4), 256–271. 10.1038/s41579-024-01116-z

8. Borsa, N., Di Pasquale, M., & Restrepo, M. I. (2019). Animal Models of Pneumococcal pneumonia. International Journal of Molecular Sciences, 20(17), 4220. 10.3390/ijms20174220

9. Chiavolini, D., Pozzi, G., & Ricci, S. (2008). Animal Models of *Streptococcus pneumoniae* Disease. Clinical Microbiology Reviews, 21(4), 666–685. 10.1128/CMR.00012-08

10. Watson, M. E., Neely, M. N., & Caparon, M. G. (n.d.). Animal Models of Streptococcus pyogenes Infection.

11. Ikuta, K. S., Swetschinski, L. R., Robles Aguilar, G., Sharara, F., Mestrovic, T., Gray, A. P., Davis Weaver, N., Wool, E. E., Han, C., Gershberg Hayoon, A., Aali, A., Abate, S. M., Abbasi-Kangevari, M., Abbasi-Kangevari, Z., Abd-Elsalam, S., Abebe, G., Abedi, A., Abhari, A. P., Abidi, H., … Naghavi, M. (2022). Global mortality associated with 33 bacterial pathogens in 2019: A systematic analysis for the Global Burden of Disease Study 2019. The Lancet, 400(10369), 2221–2248. 10.1016/S0140-6736(22)02185-7

12. Yusoff, M., Yeo, C. C., Nasir, M. H., & Amonov, M. (2024). Endonuclease A in Streptococcus pneumoniae: Escaping from neutrophil extracellular traps (NETs) and relationship in immunogenicity. Exploration of Medicine, 626–640. 10.37349/emed.2024.00245

13. Brooks, L. R. K., & Mias, G. I. (2018). Streptococcus pneumoniae’s Virulence and Host Immunity: Aging, Diagnostics, and Prevention. Frontiers in Immunology, 9, 1366. 10.3389/fimmu.2018.01366

14. Daigle, D., Morales, G., Hayford, K., Haridy, H., Abalos, M. G., Taysi, B. N., & Fletcher, M. A. (2025). Recommendations for pneumococcal vaccination of at-risk children: A global overview (2022–2024). Vaccine, 63, 127669. 10.1016/j.vaccine.2025.127669

15. King, L. M., Andrejko, K. L., Kobayashi, M., Xing, W., Cohen, A. L., Self, W. H., Resser, J. J., Whitney, C. G., Baughman, A., Kio, M., Grijalva, C. G., Traenkner, J., Rouphael, N., & Lewnard, J. A. (2024). Pneumococcal serotype distribution and coverage of existing and pipeline pneumococcal vaccines. 10.1101/2024.12.12.24318944

16. Ralph, A. P., & Carapetis, J. R. (2012). Group A Streptococcal Diseases and Their Global Burden. In G. S. Chhatwal (Ed.), Host-Pathogen Interactions in Streptococcal Diseases (Vol. 368, pp. 1–27). Springer Berlin Heidelberg. 10.1007/82_2012_280

17. Thacharodi, A., Hassan, S., Vithlani, A., Ahmed, T., Kavish, S., Geli Blacknell, N.-M., Alqahtani, A., & Pugazhendhi, A. (2025). The burden of group A Streptococcus (GAS) infections: The challenge continues in the twenty-first century. iScience, 28(1), 111677. 10.1016/j.isci.2024.111677

18. Neumann, A., Happonen, L., Karlsson, C., Bahnan, W., Frick, I.-M., & Björck, L. (2021). Streptococcal protein SIC activates monocytes and induces inflammation. iScience, 24(4), 102339. 10.1016/j.isci.2021.102339

19. Fernie-King, B. A., Seilly, D. J., Willers, C., Würzner, R., Davies, A., & Lachmann, P. J. (2001). Streptococcal inhibitor of complement (SIC) inhibits the membrane attack complex by preventing uptake of C567 onto cell membranes. Immunology, 103(3), 390–398. 10.1046/j.1365-2567.2001.01249.x

20. Åkesson, P., Herwald, H., Rasmussen, M., HÅkansson, K., Abrahamson, M., Hasan, A. A. K., Schmaier, A. H., Müller-Esterl, W., & Björck, L. (2010). Streptococcal inhibitor of complement-mediated lysis (SIC): An anti-inflammatory virulence determinant. Microbiology, 156(12), 3660–3668. 10.1099/mic.0.039578-0

21. A. Bennig, I., Ströbaek, J., Mamede, R., Sellerberg, A., Neumann, A., Friães, A., Ramirez, M., Hall, M., Collin, M., Malmström, L., Ekström, S., Frick, I.-M., Björck, L., & Happonen, L. J. (2026). Molecular mechanism of complement interference by Streptococcus pyogenes nuclease A. Communications Biology, 9(1), 1245. 10.1038/s42003-026-10956-9

22. Aghababa, H., Ting, Y. T., Pilapitiya, D., Loh, J. M. S., Young, P. G., & Proft, T. (2022). Complement evasion factor (CEF), a novel immune evasion factor of *Streptococcus pyogenes*. Virulence, 13(1), 225–240. 10.1080/21505594.2022.2027629

23. Tang, D., Khakzad, H., Hjortswang, E., Malmström, L., Ekström, S., Happonen, L., & Malmström, J. (2024). Streptolysin O accelerates the conversion of plasminogen to plasmin. Nature Communications, 15(1), 10212. 10.1038/s41467-024-54173-6

24. Zhu, L., Olsen, R. J., Lee, J. D., Porter, A. R., DeLeo, F. R., & Musser, J. M. (2017). Contribution of Secreted NADase and Streptolysin O to the Pathogenesis of Epidemic Serotype M1 Streptococcus pyogenes Infections. The American Journal of Pathology, 187(3), 605–613. 10.1016/j.ajpath.2016.11.003

25. Wenig, K., Chatwell, L., Von Pawel-Rammingen, U., Björck, L., Huber, R., & Sondermann, P. (2004). Structure of the streptococcal endopeptidase IdeS, a cysteine proteinase with strict specificity for IgG. Proceedings of the National Academy of Sciences, 101(50), 17371–17376. 10.1073/pnas.0407965101

26. Von Pawel-Rammingen, U., & Bjorck, L. (2003). IdeS and SpeB: Immunoglobulin-degrading cysteine proteinases of. Current Opinion in Microbiology, 6(1), 50–55. 10.1016/S1369-5274(03)00003-1

27. Sekuloski, S., Batzloff, M. R., Griffin, P., Parsonage, W., Elliott, S., Hartas, J., O’Rourke, P., Marquart, L., Pandey, M., Rubin, F. A., Carapetis, J., McCarthy, J., & Good, M. F. (2018). Evaluation of safety and immunogenicity of a group A streptococcus vaccine candidate (MJ8VAX) in a randomized clinical trial. PLOS ONE, 13(7), e0198658. 10.1371/journal.pone.0198658

28. Gao, N. J., Uchiyama, S., Pill, L., Dahesh, S., Olson, J., Bautista, L., Maroju, S., Berges, A., Liu, J. Z., Zurich, R. H., Van Sorge, N. M., Fairman, J., Kapoor, N., & Nizet, V. (2021). Site-Specific Conjugation of Cell Wall Polyrhamnose to Protein SpyAD Envisioning a Safe Universal Group A Streptococcal Vaccine. Infectious Microbes and Diseases, 3(2), 87–100. 10.1097/IM9.0000000000000044

29. Bensi, G., Mora, M., Tuscano, G., Biagini, M., Chiarot, E., Bombaci, M., Manetti, A. G. O., Donato, P., Swennen, E., Gallotta, M., Garibaldi, M., Pinto, V., Chiappini, N., Musser, J. M., Janulczyk, R., Mariani, M., Scarselli, M., Telford, J. L., Grifantini, R., … Grandi, G. (n.d.). Multi High-Throughput Approach for Highly Selective Identification of Vaccine Candidates: The Group A Streptococcus Case*□S.

30. Aziz, R. K., Ismail, S. A., Park, H., & Kotb, M. (2004). Post-proteomic identification of a novel phage-encoded streptodornase, Sda1, in invasive M1T1 *Streptococcus pyogenes*. Molecular Microbiology, 54(1), 184–197. 10.1111/j.1365-2958.2004.04255.x

31. Uchiyama, S., Andreoni, F., Schuepbach, R. A., Nizet, V., & Zinkernagel, A. S. (2012). DNase Sda1 Allows Invasive M1T1 Group A Streptococcus to Prevent TLR9-Dependent Recognition. PLoS Pathogens, 8(6), e1002736. 10.1371/journal.ppat.1002736

32. Zhu, L., Kuang, Z., Wilson, B. A., & Lau, G. W. (2013). Competence-Independent Activity of Pneumococcal Enda Mediates Degradation of Extracellular DNA and Nets and Is Important for Virulence. PLoS ONE, 8(7), e70363. 10.1371/journal.pone.0070363

33. Moon, A. F., Midon, M., Meiss, G., Pingoud, A., London, R. E., & Pedersen, L. C. (2011). Structural insights into catalytic and substrate binding mechanisms of the strategic EndA nuclease from Streptococcus pneumoniae. Nucleic Acids Research, 39(7), 2943–2953. 10.1093/nar/gkq1152

34. Frick, I.-M., Happonen, L., Wrighton, S., Nordenfelt, P., & Björck, L. (2023). IdeS, a secreted proteinase of Streptococcus pyogenes, is bound to a nuclease at the bacterial surface where it inactivates opsonizing IgG antibodies. Journal of Biological Chemistry, 299(11), 105345. 10.1016/j.jbc.2023.105345

35. Happonen, L., Hauri, S., Svensson Birkedal, G., Karlsson, C., De Neergaard, T., Khakzad, H., Nordenfelt, P., Wikström, M., Wisniewska, M., Björck, L., Malmström, L., & Malmström, J. (2019). A quantitative Streptococcus pyogenes–human protein–protein interaction map reveals localization of opsonizing antibodies. Nature Communications, 10(1), 2727. 10.1038/s41467-019-10583-5

36. Moon, A. F., Krahn, J. M., Lu, X., Cuneo, M. J., & Pedersen, L. C. (2016). Structural characterization of the virulence factor Sda1 nuclease from *Streptococcus pyogenes*. Nucleic Acids Research, 44(8), 3946–3957. 10.1093/nar/gkw143

37. Hughes, C. S., Foehr, S., Garfield, D. A., Furlong, E. E., Steinmetz, L. M., & Krijgsveld, J. (2014). Ultrasensitive proteome analysis using paramagnetic bead technology. Molecular Systems Biology, 10(10), MSB145625. 10.15252/msb.20145625

38. Trevor, C. E., Gonzalez-Munoz, A. L., Macleod, O. J. S., Woodcock, P. G., Rust, S., Vaughan, T. J., Garman, E. F., Minter, R., Carrington, M., & Higgins, M. K. (2019). Structure of the trypanosome transferrin receptor reveals mechanisms of ligand recognition and immune evasion. Nature Microbiology, 4(12), 2074–2081. 10.1038/s41564-019-0589-0

39. Calmettes, C., Alcantara, J., Yu, R.-H., Schryvers, A. B., & Moraes, T. F. (2012). The structural basis of transferrin sequestration by transferrin-binding protein B. Nature Structural & Molecular Biology, 19(3), 358–360. 10.1038/nsmb.2251

40. Noinaj, N., Easley, N. C., Oke, M., Mizuno, N., Gumbart, J., Boura, E., Steere, A. N., Zak, O., Aisen, P., Tajkhorshid, E., Evans, R. W., Gorringe, A. R., Mason, A. B., Steven, A. C., & Buchanan, S. K. (2012). Structural basis for iron piracy by pathogenic Neisseria. Nature, 483(7387), 53–58. 10.1038/nature10823

41. Eichenbaum, Z., Muller, E., Morse, S. A., & Scott, J. R. (1996). Acquisition of iron from host proteins by the group A streptococcus. Infection and Immunity, 64(12), 5428–5429. 10.1128/iai.64.12.5428-5429.1996

42. Ge, R., & Sun, X. (2014). Iron acquisition and regulation systems in Streptococcus species. Metallomics, 6(5), 996. 10.1039/c4mt00011k

43. Bhakdi, S., & Tranum-Jensen, J. (1982). Terminal membrane C5b-9 complex of human complement: Transition from an amphiphilic to a hydrophilic state through binding of the S protein from serum. The Journal of Cell Biology, 94(3), 755–759. 10.1083/jcb.94.3.755

44. Murphy, B. F., Kirszbaum, L., Walker, I. D., & d’Apice, A. J. (1988). SP-40,40, a newly identified normal human serum protein found in the SC5b-9 complex of complement and in the immune deposits in glomerulonephritis. Journal of Clinical Investigation, 81(6), 1858–1864. 10.1172/JCI113531

45. Mortensen, S. A., Sander, B., Jensen, R. K., Pedersen, J. S., Golas, M. M., Jensenius, J. C., Hansen, A. G., Thiel, S., & Andersen, G. R. (2017). Structure and activation of C1, the complex initiating the classical pathway of the complement cascade. Proceedings of the National Academy of Sciences, 114(5), 986–991. 10.1073/pnas.1616998114

